# Internet of Things Architecture for Cellular Biology

**DOI:** 10.1101/2021.07.29.453595

**Authors:** David F. Parks, Kateryna Voitiuk, Jinghui Geng, Matthew A. T. Elliott, Matthew G. Keefe, Erik A. Jung, Ash Robbins, Pierre V. Baudin, Victoria T. Ly, Nico Hawthorne, Dylan Yong, Sebastian E. Sanso, Nick Rezaee, Jess Sevetson, Spencer T. Seiler, Rob Currie, Alex A. Pollen, Keith B. Hengen, Tomasz J. Nowakowski, Mohammed A. Mostajo-Radji, Sofie R. Salama, Mircea Teodorescu, David Haussler

## Abstract

The Internet of Things (IoT) provides a simple framework to easily control online devices. IoT is now a commonplace tool used by technology companies, but it is rarely used in biology experiments. IoT can benefit research through alarm notifications, automation, and the real-time monitoring of experiments. We developed and implemented an IoT architecture to control biological devices used in experiments. We developed our own electrophysiology, microscopy, and microfluidic devices so that may be controlled through a unified IoT architecture. The system allows each device to be monitored and controlled through an online web tool. We present our IoT architecture so other labs may replicate it for their own experiments.

## 1. Introduction

Cloud biology uses internet protocols to connect biological devices online. This allows live experiments to be monitored and controlled through a web application. In the past, cloud biology has been used for biology education Hossain et al. (2018, 2016), ecology Guo et al. (2015), agriculture Friha et al. (2021) and marine biology Xu et al. (2019). Cloud systems are advantageous for research experiments where live sensors are spread across vast distances. For example, ecology and marine biology experiments use cloud biology to control a fleet of sensors as they traverse through their ecosystem. Cloud biology has been suggested for the online control of high-throughput cellular biology Wong et al. (2018). A backbone of many cloud biology systems are small inexpensive computing devices that are managed by a centralized server, which are used to control aspects of a biological experiment. In particular, Raspberry Pi computers have become a standard device in many cloud biology experiments Jolles (2021).

The Internet of Things (IoT) is a framework of communication that is often used to manage multiple small devices so that they are able to work in unison. IoT has become commonplace as technology used in home sensors, distributed robotic factories, and personal wearables. The framework is designed for devices to be easily connected together and controlled through underlying messaging protocols like MQTT (Message Queuing Telemetry Transport). IoT has been less commonly used for cloud biology, however, examples exist from ecology and Amazon Alexa integration of lab devices Knight et al. (2020), to commercial devices Perkel (2017a). To our knowledge, no IoT system has yet been developed for cellular biology.

IoT systems can provide many benefits to cloud-based biology experiments. IoT provides a standardized framework of communication that dramatically reduces the amount of effort required to connect each device to the cloud. Fleets of devices can be controlled with negligibly more effort than that of controlling a single device, because of the modular nature of the IoT framework. Live data streaming becomes possible using the same straightforward protocols as basic communication. IoT also provides its own method for instant notifications. This is particularly useful when an alarm notification should be sent to a scientist notifying them that their experiment is in danger Perkel (2017b).

In this article, we introduce an IoT architecture for cell biology. We demonstrate the architecture and its usage with laboratory benchtop experiments in electrophysiology, microscopy, and microfluidics. The electrophysiology, microscopy, and microfluidics devices were all built by us. They all use a standardized Raspberry Pi IoT architecture so that the system is unified and simple to control. Our IoT system provides us with real-time control and monitoring of live experiments through an online web tool. This provides us with the ability to automate research and receive live updates of the health of experiments. This architecture benefits our research and will hopefully benefit other labs who implement something similar.

## 2. System Design

Cost, scalability, maintainability, and scientific reproducibility were the fundamental requirements for our high throughput experimentation software. Low-cost is made possible by cloud computing platforms offering affordable commodity compute and storage resources at supercomputer scales. Scalability and maintainability are achieved through IoT management of devices and software containerization of data analysis processes, which both offer plug-and-play approaches with minimal dependencies between components. Scientific reproducibility is embedded through standards-based workflow definitions using Nextflow and Dockstore.

Figure 1 depicts the high-level overview of the system. Data acquisition modules (devices) execute experiments in the lab. Each module performs a specific task such as electrophysiology, microscopy, and biochemical assays. Users interact with the devices through a web-based user interface, or a lower-level software API. The software API controls devices and enables any program to control the flow of experiments. Logistics of device management, communication, and data storage are handled through the Pacific Research Platform (PRP, a nonprofit) and Amazon Web Services (AWS, for profit). In the following sections, we describe each component of the architecture.

**Figure 1:**
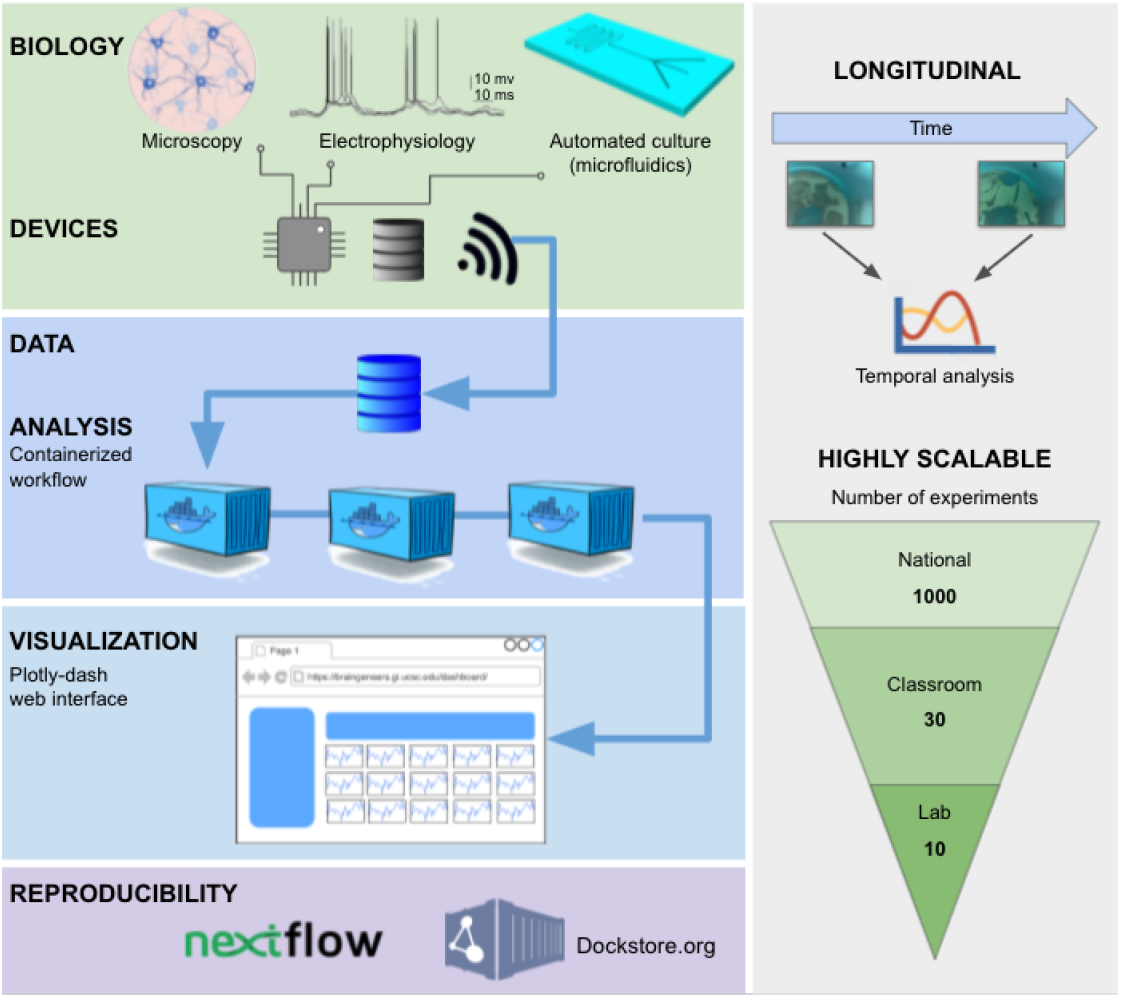
IoT Cloud Laboratory: Experiments are automated through cloud connected devices to allow scalability, reproducibility, and online monitoring.

### 2.1. Device management, communication, and control using IoT and MQTT

The data acquisition modules are lightweight and general-purpose IoT devices. The IoT devices connect to the various services that support user control, data storage, analysis, and visualization via the MQTT (Message Queuing Telemetry Transport) protocol. MQTT is a well-supported, industry-standard publish-subscribe messaging protocol.

Figure 2 depicts the central role MQTT plays in coordinating data acquisition modules and user interface communications. The MQTT protocol maintains the state and connection status for each device. It also provides a simple, lightweight publish-subscribe platform with defined *topics*. The *topics* are used by devices or user interface components to organize communication. There are two types of *topics*: a *topic* per each device (electrophysiology, microscopy, or any device performing experimental measurements or recording), and a *topic* per each running experiment. Each experiment is also assigned a UUID (Universally Unique IDentifier) which becomes an *active topic* for the period of operation.

**Figure 2:**
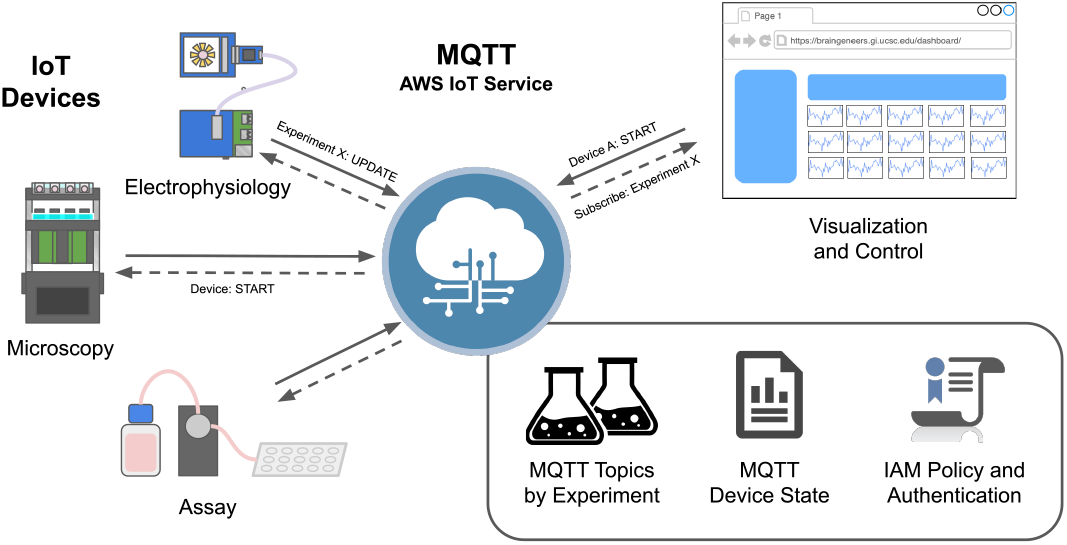
Inter-device MQTT message broker. The MQTT message broker provides integration and control over multiple internet-connected instruments. The functionality supports *clients*, data acquisition modules or software applications, to connect and subscribe to topics set by a *publisher*, such as the user interface (UI), with the proper authentication protocols. By doing so, *clients* subscribed to the topic will be informed of the state of each data acquisition module (e.g., start, stop, etc.) and parameter changes throughout an experiment.

An experiment starts when MQTT messages are published to the appropriate experiment and device *topics*. Devices subscribed to those *topics* receive the messages and take the appropriate action. Actions can also be taken automatically based on sensor readings. For example, a temperature sensor that detects overheating can publish an emergency stop message to the appropriate devices, and turn this device off. Actions may involve sending users alerts explaining errors or requesting intervention.

### 2.2. Data storage using Ceph/S3

Figure 3 shows how devices store experimental data. Primary storage and data processing are implemented on the PRP through a distributed commodity compute cluster based on Kubernetes and the Ceph (Weil et al., 2006) distributed file system. Ceph provides a highly scalable S3 interface to a virtually unlimited data store. Ceph/S3 is the primary storage for all datasets, small to terabyte-scale, commonly recorded by electrophysiology, microscopy, and biochemical assays. Our larger parallelized data processing tasks have peaked at over 5 GB/sec of concurrent I/O from S3, demonstrating the substantial scalability of the file system. Access to the Ceph/S3 data store is universally available on the internet, making it an excellent place to share large datasets across institutions.

**Figure 3:**
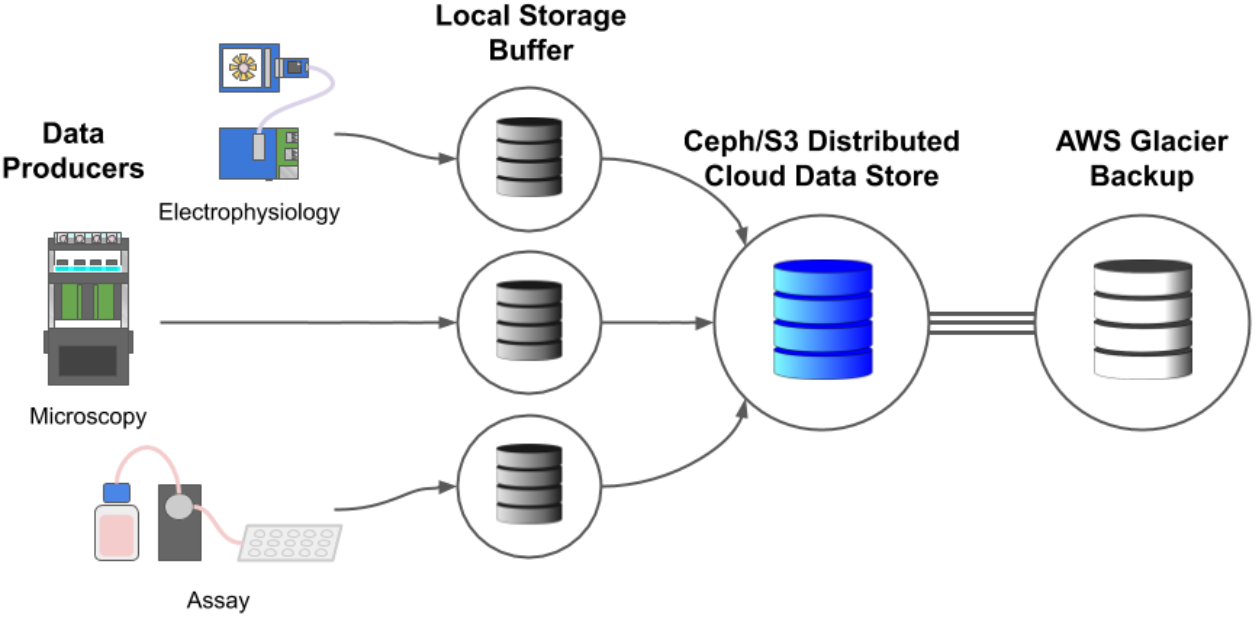
Data storage architecture. Data storage is buffered to the local device before being delivered to cloud S3 storage. Network and cloud service disruptions are expected. With the real-time data feed, interruptions only impact active visualizations of the data, which is acceptable, but the loss of experimental data is not. Each device buffers data to its local storage before making a best-effort attempt to upload it to the S3 distributed object store. Data may be buffered until the local storage is exhausted (typically enough for at least a day). The S3 distributed store is backed up to AWS Glacier to guard against user error (accidental deletion) and the loss of the S3 service. Cloud providers like AWS, GCP, and Azure have strong S3 service level agreements, unlike academic clusters such as the PRP.

As a research-oriented compute cluster, the PRP (Pacific Research Platform) does not provide strong SLAs (Service Level Agreements) for the data store. Network outages due to local network, power, or user error can cause temporary service disruptions. No guarantee is made against data loss, though the Ceph filesystem provides mechanisms to guard against common failures such as losing a node or storage media. We mitigate against data loss by scheduling a Kubernetes Cron Job with a nightly backup of all data from Ceph/S3 to AWS Deep Glacier, a cloud IaaS (Infrastructure as a Service) service providing a long-term tape storage solution. Also, all data-producing edge devices maintain a local cache that can withstand a temporary service disruption.

### 2.3. User interface using Plotly Dash

A Plotly Dash^3^ interface is easy to develop and code in Python, a common language for data science. Plotly offers a rich set of interactive plotting functionality, including specialized biology-focused visualizations. Dash provides a template to build user interfaces that implement the Observer Design Pattern (Gamma, 1995) making for an extensible and maintainable environment.

A dedicated server runs a single Plotly Dash instance under which the user interface and visualizations run as a multi-page Plotly Dash web application (see the “Visualization and control” in Figure 4). This topic will be further discussed in the “Results and Discussion” section (Figure 6 “Control’). This application can plot data from past experiments saved on Ceph/S3, or it can publish MQTT messages to the device or experiment *topics* in real time. Figure 6 and Section 3 shows a how a user would visualize a “Piphys” electrophysiology device streaming data.

**Figure 4:**
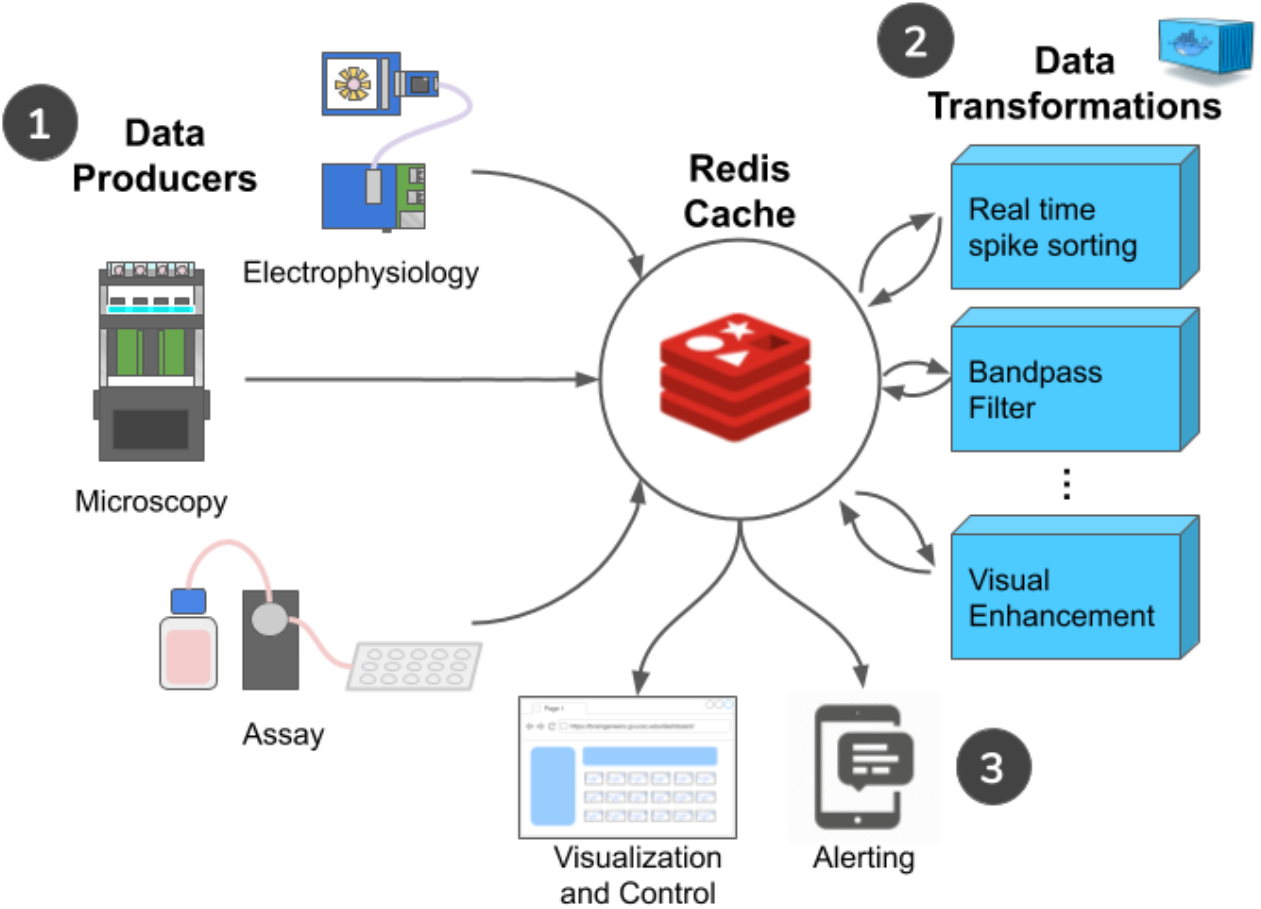
Real-time data visualization: (1) Electrophysiology, Microscopy, and Experimental Assay IoT devices produce real-time data streams on-demand only when a user is connected to a visualization that utilizes that stream. (2) Data transformations process raw data into a variety of helpful forms. Each independently containerized transformation reads a data stream and produces a new data stream. (3) Visualization and alerting notify IoT devices via MQTT that data streams are needed.

**Figure 5:**
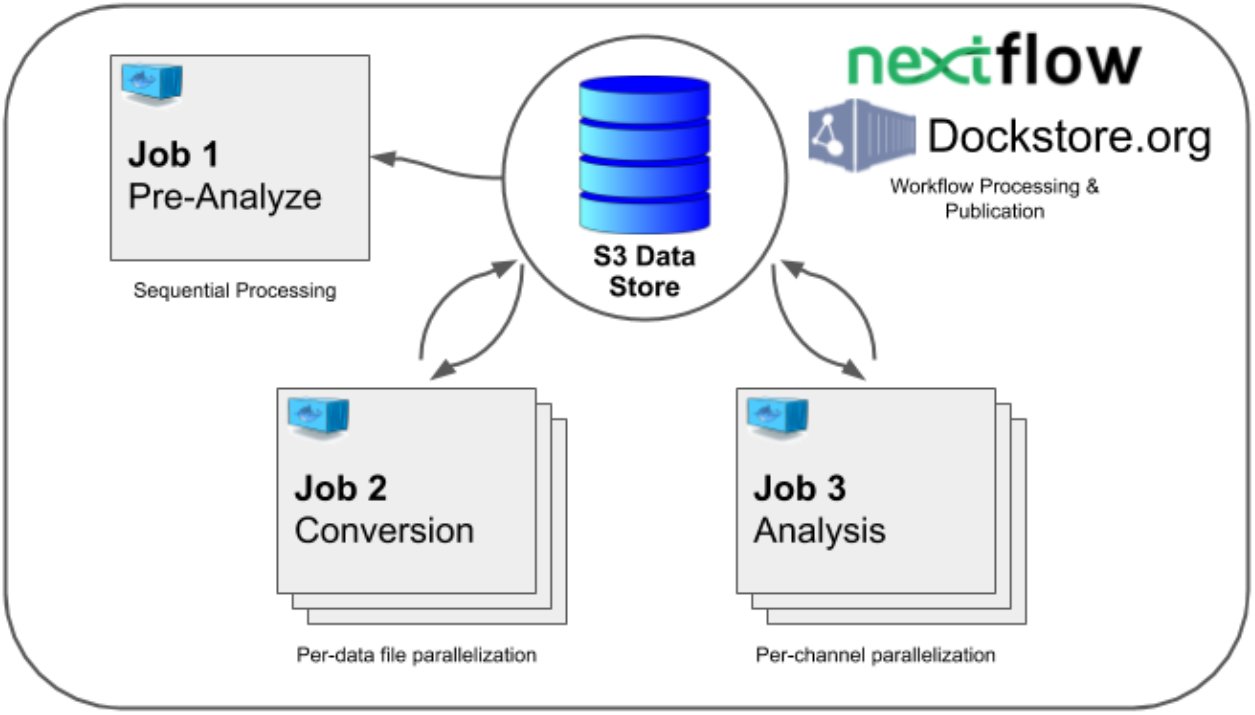
An electrophysiology experiment post data processing workflow. (Job 1) A subset of the data is analyzed to determine which channels are active, (Job 2) raw data for each active channel is converted into the form necessary for data analysis (this step takes advantage of cluster parallelism, splitting tasks by data file), and (Job 3) the data analysis including spike sorting, and other custom analysis tasks, is performed in parallel by an active channel.

**Figure 6:**
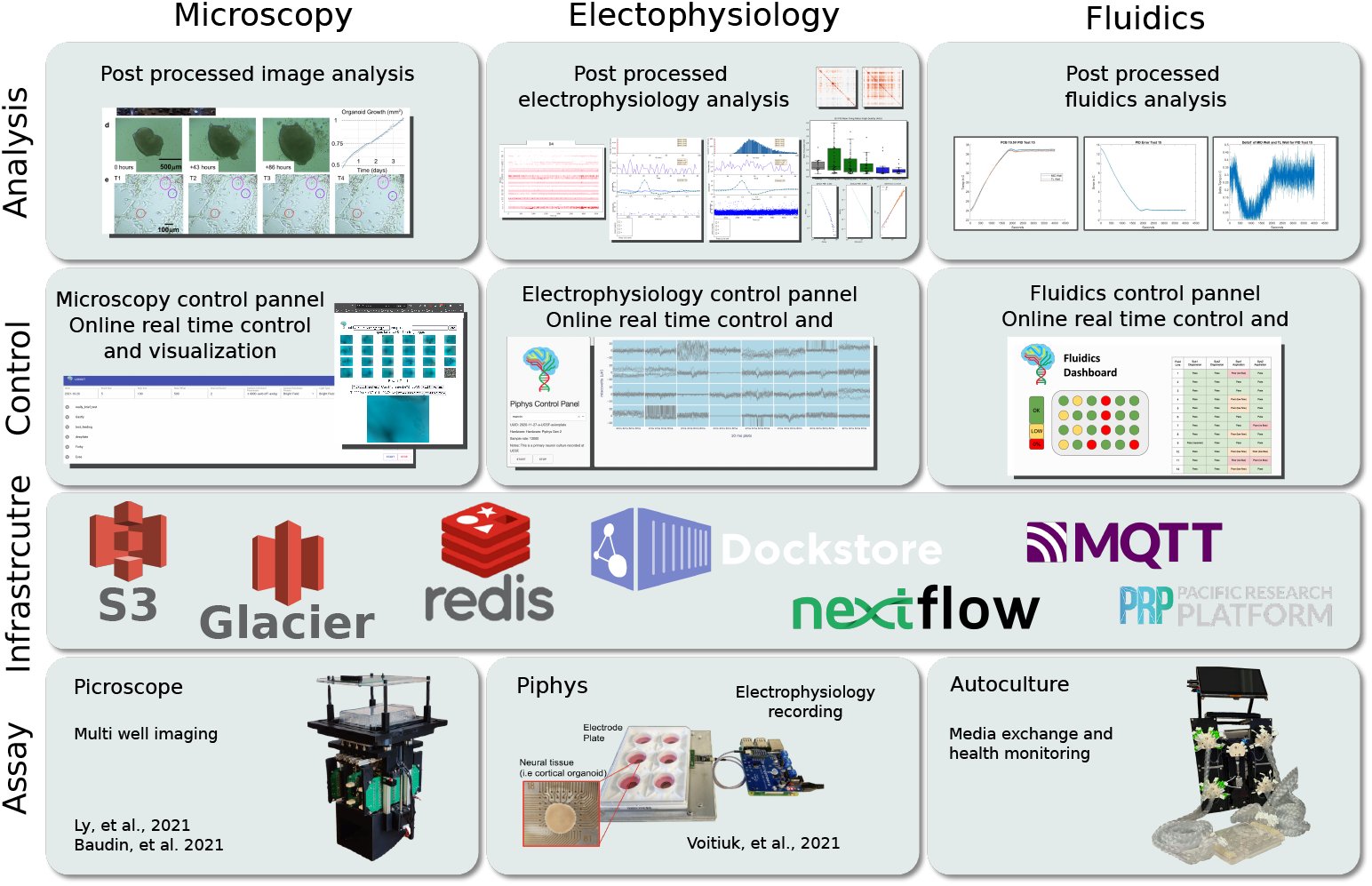
An outline of existing tools that utilize the platform described in this paper. (Assay) shows the Picroscope Ly et al. (2021); Baudin et al. (2021) for microscopy, the Piphys Voitiuk et al. (2021) for electrophysiology recording, and a fully automated auto-culture system for media recording and replacement. (Infrastructure) shows the primary suite of tools introduced in section 2.2, 2.4 and 2.5 in this paper. (Control) shows a snapshot of existing web based control of the edge devices. These web pages are running on a server in UCSC Genomics Institute. (Analysis) demos some of the reports produced as a result of the workflows that run as post-processing jobs. (the “Picroscope” and “Piphys” figures are adapted from Ly et al. (2021); Baudin et al. (2021) and Voitiuk et al. (2021))

### 2.4. Data streaming using Redis

Real-time streaming and real-time feedback are facilitated through a Redis service. Redis is a high speed database that acts as an inter-server and inter-process communication service. It is straightforward to interact with Redis using many languages, including Bash, Python, and C. Raw data feeds are sent to Redis only when the user is actively interacting with a data stream, e.g., looking at a real-time visualization, the UI client sends MQTT keep-alive messages to keep the data stream active. While MQTT is appropriate for small messages, Redis is the main communication method for larger blocks of data.

Figure 4 introduces a mechanism for handling large-scale real-time data streams. Redis provides common data structures with the inter-process locking required to coordinate between services running on separate devices. It provides a way for a data producer to publish a real-time stream of data, such as an electrophysiology recording, and for a consumer of that data, such as the Plotly Dash UI, to coordinate with each other without direct dependencies between them. Data transformations using Redis shown in Figure 4 are discussed in Section 2.6. A Redis stream is effectively a queue that can be capped in length, so that old data is automatically dropped once the maximum size of the stream is reached. Consumers, such as the Plotly Dash website, can send a recurring MQTT message to the relevant data producer to start the data stream and read the data as it is produced. A Redis service interruption merely pauses data visualization. The data producers stream a raw data feed to Redis in real-time while logging data in batches to Ceph/S3. The Ceph/S3 object store remains the primary source for data storage, and the data transfer to Ceph/S3 is resilient to service disruptions. There is no guarantee against the loss of data in the streaming approach, which is why Ceph/S3 is the primary datastore, and the Redis stream is reserved for visualizations that can incur service interruptions without lasting consequences.

### 2.5. Data processing using containerization and workflow definitions

Longitudinal electrophysiology, microscopy and microfluidic experiments commonly produce datasets on the multi-terabyte scale. Big data analysis is performed using containerized workflows built with Docker and Kubernetes and then deployed using Nextflow. Large scale machine learning especially relies on S3 for reading terabyte scale datasets. Data analytics tasks such as neural voltage signal analysis, machine learning, and image analysis require substantial computing resources and processing in multiple stages.

To provide substantial computation power and resources with simple cloud management, we utilize containerization in our infrastructure. This is the process of packaging up code and all its dependencies, so an application runs reliably in any computing environment because the code and all of its software dependencies, APIs, and versions are packaged into a binary. Containers are efficient and lightweight, they share a single host operating system (OS), and each container acts as an independent virtual machine without additional overhead (unlike full hypervisor virtual machines which replicate the OS). The container can be uploaded to a repository (for example, on Docker Hub) and downloaded and run on any computer. This includes servers in a cluster or a local lab computer.

We introduced Dockstore.org (O’Connor et al., 2017) in our design as the next logical step in scientific reproducibility, building on containerization technology. Dockstore.org is a website dedicated to hosting containerized scientific workflows formalized by workflow definitions. The formal definition of a workflow is its inputs, outputs, steps, dependencies, and the containers they run on. A common workflow language formalizes a containerized software process to ensure that organizations can run each other’s software in a standards-compliant manner. Several formal workflow definition languages exist: Nextflow (Di Tommaso et al., 2017), Common Workflow Language (CWL) (Amstutz et al., 2016), and Workflow Description Language (WDL) and are all supported by Dockstore.

Besides being a formalized workflow language, Nextflow provides a workflow runtime engine capable of deploying containerized processes to various platforms such as Kubernetes, AWS, Google Cloud, and Azure. Figure 5 depicts a standard electrophysiology data processing workflow we developed run by Nextflow and deployed to the Kubernetes-based platform on the PRP. All workflows receive a standard UUID (Universally Unique IDentifier) pointer to a dataset, allowing the workflows to find the raw data or pre-processed data produced by a dependent workflow.

#### 2.5.1. Example: electrophysiology data processing workflow

Let us consider an example workflow for an electrophysiology experiment. The goal is to detect the action potentials (spikes) of neurons by analyzing voltage recordings on multiple channels. This is part of a larger procedure called “spike sorting”. The workflow consists of 3 Jobs that occur in sequence:

For Job 1, a subset of the electrophysiology data is scanned to identify active channels. A JSON file with active channel information is recorded to Ceph/S3. This step requires a single task/container to run.

For Job 2, the dataset is converted from its raw 2-byte data (`int16`) format into a 4-byte floating-point format (`float32`) necessary for data analysis. Since the dataset is typically large (commonly in Terabytes), the information is stored in multiple files where the number of files is directly proportional to the size of the deadset. The substantial compute and I/O workload must be distributed across the cluster. One job per data file is launched, downloading the raw 2-byte data file from Ceph/S3 and uploading a 4-byte data file (`float32`) back to a temporary location on Ceph/S3. In the process, the data is separated into individual channels for processing in the next step. Notice that the conversion process must download and re-upload the full dataset because multi-terabyte datasets are too big to fit on local nodes.

For Job 3, the data for active channels is pulled from the distributed filesystem, then spike sorting and spike timing analysis is performed. The results are placed back on the distributed filesystem.

While each job runs in series and depends on the last, there are no dependencies between the jobs other than those involving the data that is posted to the primary datastore Ceph/S3. Each dataset has a unique ID (UUID) which also serves as a location pointer to where data is stored on the Ceph/S3. This UUID is the only parameter passed between jobs.

Beside the illustrated example in this section, Figure 7 shows a more general overview of resources employed and parallelization of the data processing by workflows including imaging and biochemical assays.

**Figure 7:**
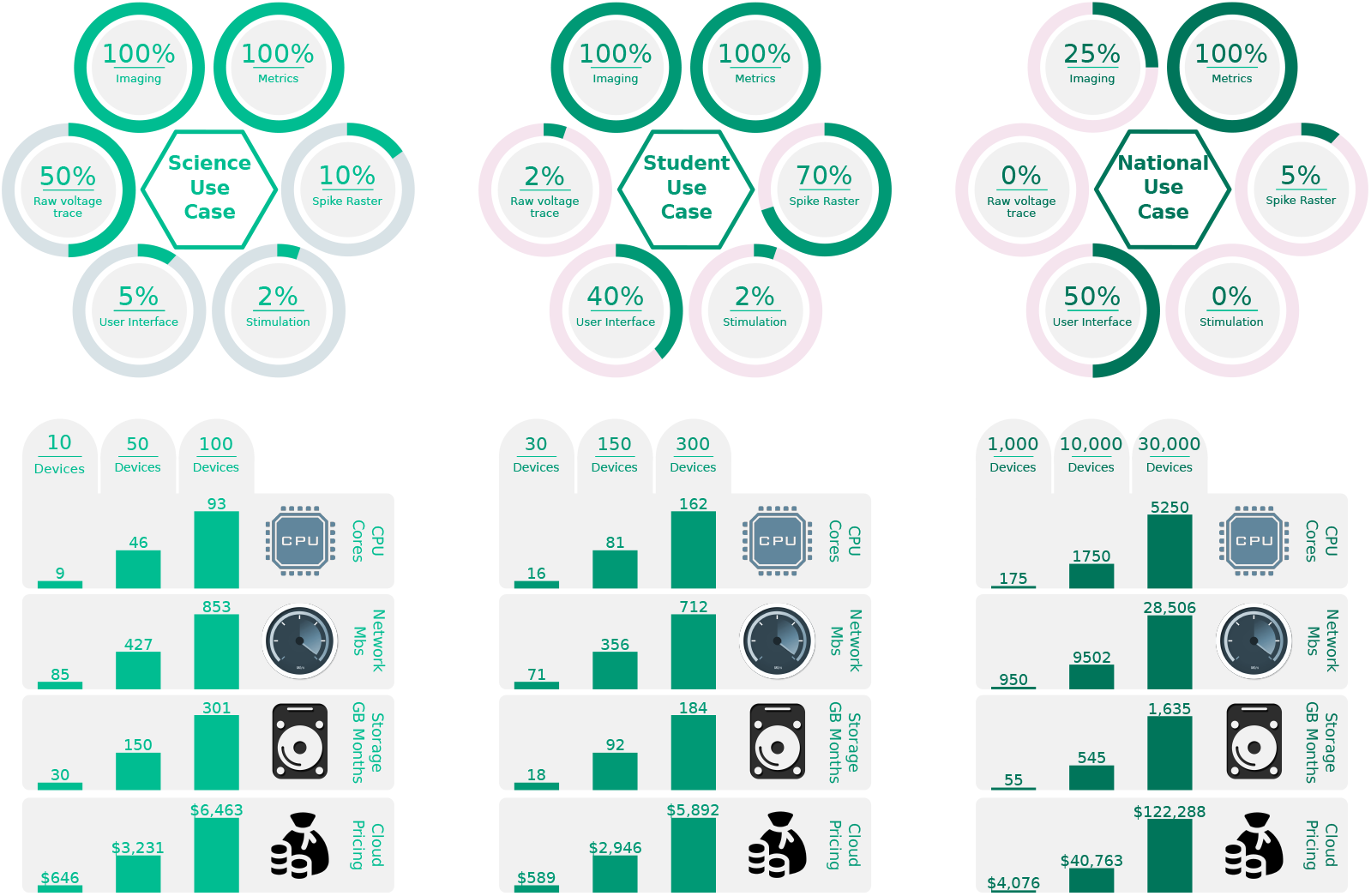
Monthly resource utilization requirements given three use cases: Science, Student, and National scales. The assumed distribution of device functions under each use case is displayed above. Resource utilization for CPU, Network, and Storage are displayed below. An estimate of cloud compute costs are provided on the bottom. The number of active devices varies from fewer in the Science use to many in the National use. We define “% imaging,” as the percentage of devices actively recording and storing microscopy images; “% metrics,” as the percentage of devices actively recording measurements such as media concentrations and temperatures; “% raw voltage,” as the percentage of devices recording and storing full raw voltage traces across all electrophysiology channels; “% spike raster,” as the percentage of devices registering only neural spikes events (estimated to be 10% of the raw voltage data); “% UI,” as the number of active users on the web interface relative to the total number of devices; and “% stimulation,” as the percentage of devices that are actively executing electrode stimulation requests.

### 2.6. Real-time analysis, data processing, and transformations

Deploying containerized workflows via Nextflow works well for large-scale post-processing and data analysis but does not provide a mechanism for real-time visualizations and experiment control.

The Redis in-memory database service coordinates the real-time exchange of data between many producers and consumers. For example, an electrophysiology recording on 32 channels at 25 kHz will produce a data stream of 1.6 MB/sec, which a user may want to monitor in real-time. Equivalently, a microscopy recording could provide a stream of images for visualization for real-time experimental metrics, such as 600 MB every recording for all the images with 24 cameras each capturing 10 pictures.

Transformation of data with visual enhancements applied in real-time is often more informative than seeing raw data. Data transformations are performed by containerized processes that read a stream of data and write a new stream of transformed data. For example, a container reads a raw electrophysiology stream and writes a new steam with the bandpass filtered data. After applying the data transformation, a visualization such as a Plotly Dash web page would read the appropriate data stream output. Data transformations have no dependencies other than the Redis stream they read from and can be entirely independent workflows. Transformations can easily be added or changed without changing any other software infrastructure components.

## 3. Results and Discussion

This software architecture supports different modes of data acquisition that measure and report data. Here we focus on three types of modules for proof of concept: (1) Electrophysiology – recording and stimulation of neural cell cultures (2) Microscopy – imaging of cell cultures (3) Automated culture – feeding cells and sampling media for metabolites and RNA expression using a programmable microfluidics system. These modules are implemented and presented in Figure 6.

We will look at each of these data acquisition modules (IoT-based edge devices) in turn and discuss how they interact with the software architecture and user. For this example, we assume users will interact with devices through the web UI application. Users can be located anywhere on the Internet without concern for the location of these physical devices. This facilitates cross-campus and cross-institutional collaborations. For instance, we often perform electrophysiology and microscopy experiments from Santa Cruz on devices located 90 miles away in San Francisco. Of course, experiments still require some manipulation by a researcher at the local site (i.e., placing cell cultures on the devices and performing adjustments if components are misaligned).

To begin an electrophysiology experiment, a user opens the browser with the Plotly Dash web application (Figure 6, Control). The application queries AWS IoT service for online electrophysiology devices (Figure 6, Assay). The device can be Piphys (Voitiuk et al., 2021) or any platform/recording system whose computer runs the same code that responds to the IoT architecture and can control the system programmatically. When the user selects a device, an MQTT ‘ping’ message is sent to the relevant device every 30 seconds, indicating that a user is actively monitoring data from that device. As long as the electrophysiology device receives these pings, it will send raw data to its Redis stream (Figure 6, Infrastructure). Since the device is responsible for only a single data stream, many users can monitor and interact with the particular device without additional overhead. If the device has not received user messages for at least a minute, it will cease streaming its data. This protocol ensures the proper decoupling of users from devices, and devices are not dependent on a user gracefully shutting down the connection.

As shown in Figure 4, one or more data transformation processes can read the raw data stream and post a processed stream of data, such as real-time spike sorting. The web visualization can display the appropriate transformed data stream for the user (Figure 6, Analysis).

Stopping the experiment will automatically initiate a batch processing workflow on the Kubernetes compute platform. Users can configure the workflow to include job modules such as spike sorting, clustering, and other customized metrics of neural activity.

Microscopy, such as the Picroscope, typically operates at a lower sampling rate and over a longer continuous period than electrophysiology. Microscopy devices record images of cell culture morphology at varying focal layers and time-frequency. As with electrophysiology, these images are initially buffered locally and then flushed to the Ceph/S3 filesystem every few minutes. A user will view the data in the same web UI portal as electrophysiology. Since cell culture morphology changes relatively slowly, microscopy visualizations do not require real-time Redis streaming. The user may update the parameters of the microscopy recording with MQTT messages sent to the device *topic* updating the state.

Assay devices support the lifecycle of the cell culture, providing new media and taking regular measurements relevant to the cell culture’s health and environmental state. Much like microscopy, most of these measurements are sampled continuously over the lifetime of the culture and are posted directly to Ceph/S3 at regular time intervals. When the user accesses a UI page detailing the lifecycle of the culture, these metrics will be pulled in near real-time from Ceph/S3. The user can update and change metrics related to cells the lifecycle by an MQTT message from the UI page to the device to update its state and initiate a change in the device behavior.

### 3.1. Scaling

In the previous section, we considered one experiment with a few data acquisition modules running in a single lab. This section considers hypothetical studies of tens to thousands of experiments operating simultaneously. Each user will use different features of the devices, and there would be a virtually infinite combinations of features when many devices are deployed. We define three use cases and provide an analysis of these and their assumptions, we call the use cases, Science, Student, and National. We provide a distribution over the basic functions and devices that we expect the users will employ in each case. For each case we provide estimates of CPU, Network, and Storage resources required, visualized in Figure 7. Also provided in Figure 7 is an estimate of cloud computing and storage cost based on AWS pricing. The use of the PRP academic compute cluster precludes the majority of these costs and speaks to the value the PRP brings to academic institutions.

In the Science use case, we assume a higher degree of active imaging and electrophysiology. This use case focuses on more resource-intensive lab use in the pursuit of scientific inquiry at high detail. In this configuration, storage requirements are the most significant bottleneck, growing at tens to hundreds of GB of data per hour. We find that tens of devices are appropriate for this use case before resource utilization becomes excessive.

In the Student use case, we anticipate a limited number of universities using the devices to teach classes in cell biology on live cultures hosted at a remote lab. In this use case, we assume a scale on the order of hundreds of devices. Users in this scenario will rely heavily on visualizations, including both real-time microscopy and electrophysiology. The lab that hosts hundreds of experiments with the expectation of concurrent access will require additional network bandwidth beyond what is available in a typical lab or office. At least two Gigabit network ports and matching ISP bandwidth would be necessary to support the load. At this scale, if electrophysiology is involved, limiting data that is sent over the wire to active spiking events, rather than raw signal measurements, is imperative. This requires on-device spike detection.

Lastly, in the National use case, we consider a scaled-out fleet of thousands to tens of thousands of devices. This case assumes wide-scale adoption by laboratories or secondary education facilities across the country or world. This scale requires substantial cloud computing resources to support the load and serve microscopy images and electrophysiology data to every user. It will also require significant wet lab infrastructure at the site(s) housing the biology as well as expenses of cell culture maintenance. However, given this investment, this infrastructure can enable remote experimentation by a large and diverse population.

## 4. Conclusion

This paper outlines an IoT software architecture that supports the control and analysis of electrophysiology, microscopy, and experimental assays on cell cultures. We emphasize the benefits of having a centralized online hub where automated experiments are managed through a portal. Scientists benefit from receiving notifications on the status of their experiments and monitor its progression without perturbing samples. Our architecture is built on an open-source design with scientific reproducibility in mind. Future advances in IoT architecture for cell biology may open new possibilities to scale high-throughput experiments, which may benefit drug screens, gene knockout studies and a host of other types of experiments. We hope our architecture examples will further advance the implementation of IoT in cellular biology.

## 5. Acknowledgments

This work is supported by the Schmidt Futures Foundation SF 857 (D.H.). Research reported in this publication was also supported by the National Institute Of Mental Health of the National Institutes of Health under award number R01MH120295 (S.R.S.), the National Science Foundation under award number NSF 2034037 (M.T.), the National Defense Science and Engineering Graduate Fellowship (00002116, M.G.K.), and a gift the William K. Bowes Jr Foundation (T.J.N). K.V. was supported by grant T32HG008345 from the National Human Genome Research Institute (NHGRI), part of National Institutes of Health (NIH), USA. D.H. is a Howard Hughes Medical Institute Investigator.

Through the Pacific Research Platform, this work was supported in part by NSF awards CNS-1730158, ACI-1540112, ACI-1541349, OAC-1826967, the University of California Office of the President, and the University of California San Diego’s California Institute for Telecommunications and Information Technology/Qualcomm Institute. Thanks to CENIC for the 100 Gpbs networks.

Cell cultures used in this work are from IBSC stem cell culture facilities with identifier RRID:SCR_021353.

## Author Contributions

R.C. was the chief architect for the infrastructure design, and D.F.P made it work, creating necessary integration and cross platform communication. V.T.L, P.V.B worked on microscopy. K.V., J.G., M.G.K, A.R., N.H. worked on electrophysiology and analysis and K.B.H advised. S.T.S worked on automated IoT cell culture. D.F.P. and S.E.S. created the web user interface design. D.Y. and N.R. worked on data infrastructure, backups, and automated data processing pipelines. S.R.S, A.A.P. and T.J.N. provided cell culture. D.H., M.T., S.R.S, A.A.P. and T.J.N. supervised the team and secured funding. D.F.P., K.V., J.G., and M.A.T.E., wrote the manuscript with support from E.A.J., M.A.M.R. All authors reviewed the manuscript.

## Footnotes

3 https://plotly.com

## References

Amstutz, P., Crusoe, M. R., Tijanić, N., Chapman, B., Chilton, J., Heuer, M., Kartashov, A., Leehr, D., Ménager, H., Nedeljkovich, M., Scales, M., Soiland-Reyes, S., & Stojanovic, L. (2016). Common Workflow Language, v1.0,. URL:https://www.research.manchester.ac.uk/portal/en/publications/common-workflow-language-v10(741919f5-d0ab-4557-9763-b811e911423b).html. doi:10.6084/m9.figshare.3115156.v2. Publisher: figshare.

Baudin, P. V., Ly, V. T., Pansodtee, P., Jung, E. A., Currie, R., Hoffman, R., Willsey, H. R., Pollen, A. A., Nowakowski, T. J., Haussler, D., Mostajo-Radji, M. A., Salama, S. R., & Teodorescu, M. (2021). Low cost cloud based remote microscopy for biological sciences. Internet of Things, (p. 100454). URL:https://www.sciencedirect.com/science/article/pii/S2542660521000962. doi:10.1016/j.iot.2021.100454.

Di Tommaso, P., Chatzou, M., Floden, E. W., Barja, P. P., Palumbo, E., & Notredame, C. (2017). Nextflow enables reproducible computational workflows. Nature Biotechnology, 35, 316–319. URL: https://www.nature.com/articles/nbt.3820/. doi:10.1038/nbt.3820. Number: 4 Publisher: Nature Publishing Group.

Friha, O., Ferrag, M. A., Shu, L., Maglaras, L., & Wang, X. (2021). Internet of Things for the Future of Smart Agriculture: A Comprehensive Survey of Emerging Technologies. IEEE/CAA Journal of Automatica Sinica, 8, 718–752. doi:10.1109/JAS.2021.1003925. Conference Name: IEEE/CAA Journal of Automatica Sinica.

Gamma, E. (1995). Design patterns: elements of reusable object-oriented software. Reading, Mass.: Addison-Wesley. URL:http://archive.org/details/designpatternsel00gamm.

Guo, S., Qiang, M., Luan, X., Xu, P., He, G., Yin, X., Xi, L., Jin, X., Shao, J., Chen, X., Fang, D., & Li, B. (2015). The application of the Internet of Things to animal ecology. Integrative Zoology, 10, 572–578. URL:https://onlinelibrary.wiley.com/doi/abs/10.1111/1749-4877.12162. doi:10.1111/1749-4877.12162. _eprint:https://onlinelibrary.wiley.com/doi/pdf/10.1111/1749-4877.12162.

Hossain, Z., Bumbacher, E., Brauneis, A., Diaz, M., Saltarelli, A., Blikstein, P., & Riedel-Kruse, I. H. (2018). Design Guidelines and Empirical Case Study for Scaling Authentic Inquiry-based Science Learning via Open Online Courses and Interactive Biology Cloud Labs. International Journal of Artificial Intelligence in Education, 28, 478–507. URL:https://doi.org/10.1007/s40593-017-0150-3. doi:10.1007/s40593-017-0150-3.

Hossain, Z., Bumbacher, E. W., Chung, A. M., Kim, H., Litton, C., Walter, A. D., Pradhan, S. N., Jona, K., Blikstein, P., & Riedel-Kruse, I. H. (2016). Interactive and scalable biology cloud experimentation for scientific inquiry and education. Nature Biotechnology, 34, 1293–1298. URL:https://www.nature.com/articles/nbt.3747. doi:10.1038/nbt.3747. Number: 12 Publisher: Nature Publishing Group.

Jolles, J. W. (2021). Broad-scale applications of the Raspberry Pi: A review and guide for biologists. Methods in Ecology and Evolution, 12, 1562–1579. URL:https://onlinelibrary.wiley.com/doi/abs/10.1111/2041-210X.13652. doi:10.1111/2041-210X.13652. _eprint: https://onlinelibrary.wiley.com/doi/pdf/10.1111/2041-210X.13652.

Knight, N. J., Kanza, S., Cruickshank, D., Brocklesby, W. S., & Frey, J. G. (2020). Talk2Lab: The Smart Lab of the Future. IEEE Internet of Things Journal, 7, 8631–8640. doi:10.1109/JIOT.2020.2995323. Conference Name: IEEE Internet of Things Journal.

Ly, V. T., Baudin, P. V., Pansodtee, P., Jung, E. A., Voitiuk, K., Rosen, Y. M., Willsey, H. R., Mantalas, G. L., Seiler, S. T., Selberg, J. A., Cordero, S. A., Ross, J. M., Rolandi, M., Pollen, A. A., Nowakowski, T. J., Haussler, D., Mostajo-Radji, M. A., Salama, S. R., & Teodorescu, M. (2021). Picroscope: low-cost system for simultaneous longitudinal biological imaging. Communications Biology, 4, 1–11. URL:https://www.nature.com/articles/s42003-021-02779-7. doi:10.1038/s42003-021-02779-7. Number: 1 Publisher: Nature Publishing Group.

O’Connor, B. D., Yuen, D., Chung, V., Duncan, A. G., Liu, X. K., Patricia, J., Paten, B., Stein, L., & Ferretti, V. (2017). The Dockstore: enabling modular, community-focused sharing of Docker-based genomics tools and workflows. F1000Research, 6. URL:https://www.ncbi.nlm.nih.gov/pmc/articles/PMC5333608/. doi:10.12688/f1000research.10137.1.

Perkel, J. M. (2017a). The Internet of Things comes to the lab. Nature, 542, 125–126. URL: https://www.nature.com/articles/542125a. doi:10.1038/542125a. Bandiera_abtest: a Cg_type: Nature Research Journals Number: 7639 Primary_atype: Special Features Publisher: Nature Publishing Group Subject_term: Information technology;Technology Subject_term_id: information-technology;technology.

Perkel, J. M. (2017b). The Internet of Things comes to the lab. Nature, 542, 125–126. URL:https://www.nature.com/articles/542125a. doi:10.1038/542125a. Number: 7639 Publisher: Nature Publishing Group.

Voitiuk, K., Geng, J., Keefe, M. G., Parks, D. F., Sanso, S. E., Hawthorne, N., Freeman, D. B., Mostajo-Radji, M. A., Nowakowski, T. J., Salama, S. R., Teodorescu, M., & Haussler, D. (2021). Light-weight Electrophysiology Hardware and Software Platform for Cloud-Based Neural Recording Experiments. bioRxiv, (p. 2021.05.18.444685). URL:https://www.biorxiv.org/content/10.1101/2021.05.18.444685v2. doi:10.1101/2021.05.18.444685. Publisher: Cold Spring Harbor Laboratory Section: New Results.

Weil, S., Brandt, S. A., Miller, E. L., Long, D. D. E., & Maltzahn, C. (2006). Ceph: A Scalable, High-Performance Distributed File System. In Proceedings of the 7th Conference on Operating Systems Design and Implementation (OSDI’06).

Wong, B. G., Mancuso, C. P., Kiriakov, S., Bashor, C. J., & Khalil, A. S. (2018). Precise, automated control of conditions for high-throughput growth of yeast and bacteria with eVOLVER. Nature Biotechnology, 36, 614–623. URL:https://www.nature.com/articles/nbt.4151. doi:10.1038/nbt.4151. Number: 7 Publisher: Nature Publishing Group.

Xu, G., Shi, Y., Sun, X., & Shen, W. (2019). Internet of Things in Marine Environment Monitoring: A Review. Sensors, 19, 1711. URL:https://www.mdpi.com/1424-8220/19/7/1711. doi:10.3390/S19071711. Number: 7 Publisher: Multidisciplinary Digital Publishing Institute.

